# Electron-Activated Dissociation and Collision-Induced Dissociation Glycopeptide Fragmentation for Improved Glycoproteomics

**DOI:** 10.1101/2024.02.22.581095

**Authors:** Kyle L. Macauslane, Cassandra L. Pegg, Amanda S. Nouwens, Edward D. Kerr, Joy Seitanidou, Benjamin L. Schulz

## Abstract

Tandem mass spectrometry coupled with liquid chromatography (LC-MS/MS) has proven a versatile tool for the identification and quantification of proteins and their post-translational modifications (PTMs). Protein glycosylation is a critical PTM for the stability and biological function of many proteins, but full characterisation of site-specific glycosylation of proteins remains analytically challenging. Collision induced dissociation (CID) is the most common fragmentation method used in LC-MS/MS workflows, but loss of labile modifications render CID inappropriate for detailed characterisation of site-specific glycosylation. Electron-based dissociation (ExD) methods provide alternatives that retain intact glycopeptide fragments for unambiguous site localisation, but these methods often underperform CID due to increased reaction times and reduced efficiency. Electron activated dissociation (EAD) is another strategy for glycopeptide fragmentation. Here, we use a ZenoTOF 7600 SCIEX instrument to compare the performance of various fragmentation techniques for the analysis of a complex mixture of mammalian *O*- and *N*-glycopeptides. We found CID fragmentation identified the most glycopeptides and generally produced higher quality spectra, but EAD provided improved confidence in glycosylation site localisation. Supplementing EAD with CID fragmentation (EAciD) further increased the number and quality of glycopeptide identifications, while retaining localisation confidence. These methods will be useful for glycoproteomics workflows for either optimal glycopeptide identification or characterisation.

## Introduction

Glycosylation is a post-translational modification (PTM) of most eukaryotic secreted and membrane proteins that is critical for protein folding, stability, and mediating diverse functions [1–3], and is therefore of both physiological and pathological importance [4–7]. Protein glycosylation is chemically and biosynthetically diverse, with the most well-studied and common forms in mammalian glycoproteins being *N-* and *O-*glycosylation. As secretory and membrane polypeptides are translocated into the endoplasmic reticulum (ER) they can be co- and post-translocationally *N*-glycosylated with a Glc_3_Man_9_GlcNAc_2_ glycan at select asparagine residues [8]. The likelihood that a specific asparagine residue will be *N*-glycosylated is much higher if it is located in a sequence motif known as a glycosylation sequon (N-X-S/T; X≠P), which has high affinity for the peptide acceptor binding site of the oligosaccharyltransferase (OST) enzyme that catalyses *N*-glycosylation [9–11]. In the Golgi, proteins can be *O*-glycosylated with GalNAc at serine and threonine residues, typically in serine/threonine-rich mucin domains [12]. Both *N-* and *O-*glycans can be further processed by a biosynthetic network of glycosylhydrolases and glycosyltransferases as glycoproteins traffic through the Golgi. Not all potential sites of glycosylation are necessarily occupied, and both the glycan occupancy and structures at a site can vary due to diverse regulatory processes and are subject to the overall health of a cell or organism [13–15].

In mammals, processing of *N*-glycans gives rise to three major classes of glycan structures: oligomannose, hybrid, and complex. Each contains a common core (Man_3_GlcNAc_2_) that can also be modified with core fucose (Fuc) [16]. Oligomannose glycans feature terminal branches comprised entirely of mannose (Man), complex glycans have trimming and substitution and instead terminate in sugars such as sialic acid (*N*-acetylneuraminic acid, NeuAc), while hybrid glycans feature both substituted and unsubstituted antennae [16]. *O*-GalNAc glycans can have eight core structures built from the initial GalNAc, with terminal structures similar to those found on *N*-glycans [17].

Glycoproteomics aims to identify, characterise, and quantify all features of a glycoproteome. That is, the ultimate goal involves not only the identification of glycoproteins from complex samples, but also characterisation of both the occupancy of individual sites of glycosylation (macroheterogeneity), and of the structure and composition of any attached glycans at specific sites (microheterogeneity) [18]. Hence, the ideal glycoproteomic workflow involves the characterisation of the underlying peptide backbone, the attached glycan, and of the site(s) of modification [16]. It is for this reason intact glycopeptide analysis with LC-MS/MS is the method of choice in glycoproteomics, as only analysis of intact glycopeptides can permit a complete characterisation [19].

LC-MS/MS based glyco/proteomics uses fragmentation of precursor glyco/peptide ions to characterise the resultant product ions. The dissociation techniques used in peptide fragmentation can broadly be categorised in two groups: thermal- and radical-driven dissociation techniques. The most commonly used fragmentation method in glycoproteomics is thermally driven, beam-type collision induced dissociation (CID or alternatively HCD) [20]. In CID, collision of accelerated precursor ions with neutral gas molecules imparts the internal vibrational energy that induces bond fragmentation [21]. CID results in peptide (a-, b- and y-type ions) and glycosidic bond (B- and Y-type ions) fragmentation events [16, 22]. In particular, low mass oxonium ions arising from fragmentation of mono/disaccharides are a fundamental feature of glycopeptide fragmentation with CID that enable identification of the monosaccharides present in the attached glycan [23]. CID fragmentation results in preferential cleavage of the weakest covalent bonds within a molecule, and as such occurs preferentially at glycosidic bonds over peptide C-N (amide) bonds. Glycopeptide CID fragmentation spectra are therefore rich in information about the glycan, but relatively poor in information about the peptide [24]. Nonetheless, rapid acquisition rates allow for in-depth identification of glycopeptides in complex samples [20]. However, the lack of peptide fragment ions makes localisation of the site of glycosylation difficult [25].

Radical-driven techniques, often called “soft” dissociation methods, such as electron capture dissociation (ECD) and electron transfer dissociation (ETD), generally fall under the collective term electron-driven dissociation (ExD). These techniques use electrons to impart a radical state that induces fragmentation between the stronger N–C_⍺_ (amine) bond, resulting in preferential cleavage of the peptide backbone and allowing localisation of modifications [26–28]. ExD fragmentation results in peptide bond fragmentation to generate c- and z-type fragment ions, but little to no glycosidic bond fragmentation results [29]. Although this can complicate characterisation of the glycan due to the absence of B-, Y- and oxonium ions, intact glycopeptide fragments are retained, which can be used to unambiguously determine the site of glycosylation. ExD can also provide more complete peptide backbone characterisation, as the N-C_⍺_ fragmentation in ExD is less dependent on peptide features such as charge state and amino acid sequence than in CID [30]. However, ExD techniques are potentially limited by several disadvantages. Peptides with low precursor charge states can yield low fragment ion intensities [31]. This is especially problematic in the case of dissociation of doubly charged precursors, as electron capture necessitates the generation of at least one neutral fragment that cannot be detected [32]. ExD methods are therefore most efficient for highly charged peptide precursor ions. However, the addition of neutral glycans in glycopeptides contribute to the mass of the precursor without typically increasing the positive charge [33]. Further, in the case of sialylated glycopeptides, the addition of a negatively charged glycan can decrease the overall charge [33]. Reaction times for ExD techniques are also longer than CID, with up to hundreds of milliseconds required for each spectrum, which reduces the number of spectra that can be generated in an LC-MS/MS analysis. It is for these reasons CID has typically been favoured over ExD methods for glycopeptide analysis.

Another group of fragmentation strategies worth considering are the combinations of ExD and collisional dissociation [34]. ExD and CID are complementary techniques which enable analysis of the peptide backbone with labile structures intact and the glycan structure, respectively [29]. Combining the two (i.e. electron transfer/higher-energy collision dissociation; EThcD) provides spectra with fragment ions typical of both dissociation modes, although the duty cycle costs associated with large reaction times in ExD are still retained.

The ZenoTOF 7600 (SCIEX) allows a variant of ExD known as electron activated dissociation (EAD) [35]. One key feature of EAD is the ability to leverage several tuneable parameters for specific experimental needs. Here, we optimize these parameters to allow in-depth analysis of a complex mixture of mammalian *O*- and *N*-glycopeptides.

## Methods

### Cell Culture

Human A549 cells (purchased from the ATCC) were cultured in Dulbecco’s Modified Eagle’s Medium supplemented with 50 U/mL penicillin-streptomycin (1% *v/v*) and foetal-bovine serum (10% *v/v*). Cells were cultured at 37 °C with injection of 5% CO_2_. Cells with a passage number < 25 were used to produce secreted proteins. In brief, A549 cells were grown until exceeding 90% confluence of a T175 flask (∼2×10^7^ cells). At this point, cells were washed with chilled PBS four times to remove growth serum proteins, and then supplemented with serum-free media. Spent media was collected from the cells after 24 hrs, concentrated with 10 kDa MWCO Amicon Ultra filter units (Millipore) via centrifugation at 4,000 rcf for 30 min, and exchanged into 50 mM HEPES buffer pH 7.4. Protein concentrations were determined using the Qubit Protein Quantification Assay Kit (Thermo Fisher Scientific).

### Protein Sample Preparation

Proteins (90 µg aliquots) were denatured and reduced by addition of 2x lysis buffer to give final concentrations of 1% SDS, 50 mM Tris-HCl buffer pH 8.0, and 10 mM DTT, and incubated at 95 °C for 10 min. After cooling to room temperature, proteins were alkylated by addition of acrylamide to a final concentration of 25 mM and incubation at 30 °C with shaking at 1500 rpm for 1 h. Excess acrylamide was quenched by the addition of DTT to an additional final concentration of 5 mM. Proteins were precipitated by addition of four volumes of methanol/acetone (1:1 *v/v*) and incubation at −20 °C for 16 h. Precipitated samples were centrifuged at 18,000 rcf for 10 min to pellet proteins and the supernatant was discarded. Proteins were digested by resuspension in 50 mM ammonium bicarbonate with porcine trypsin at a 1:20 enzyme to protein ratio and incubation at 37 °C with shaking at 1500 rpm for 16 h.

### Glycopeptide Enrichment

Peptides were dried using vacuum centrifugation and resuspended in 500 µL 0.1% trifluoroacetic acid (TFA) for sample clean-up with 50 mg Sep-Pak columns (Waters). Peptides underwent HILIC enrichment for glycopeptides as previously described [36], with a 1 min incubation period following sample addition. HILIC enrichment was performed in 80% ACN using PolyHYDROXYETHYL A™, 100 Å pore diameter, 12 µm particle diameter (PolyLC) HILIC beads. Eluted samples enriched for glycopeptides were dried with vacuum centrifugation and resuspended in 0.1% formic acid for LC-MS/MS analysis.

### LC-MS/MS Analysis

Samples were analysed with a ZenoTOF 7600 (SCIEX) mass spectrometer coupled to a Acquity UPLC M-Class system (Waters). Approximately 1-2 µg of enriched glycopeptide samples were injected onto a Waters nanoEase M/Z HSS T3 C18 column (300 µm x 150 mm, 1.8 µm, 100Å). The mobile phases were A: 0.1% formic acid in water, and B: 0.1% formic acid in acetonitrile. Peptides were loaded in 5% B. The LC was held at 5% B for 30 s and peptides were eluted with a linear gradient from 5-35% B over 22 min at a flow rate of 5 µL/min. ZenoTOF source conditions included: spray voltage, 5000V; Gas 1 and Gas 2, 20 psi; curtain gas, 35 psi; CAD gas, 7; temperature, 150 °C; and column temperature, 35 °C. MS1 spectra were acquired with: mass range, 300-2250 *m/z*; accumulation time, 0.2 s; Declustering Potential, 80 V; Collision Energy, 10 V; and time bins to sum, 8. The 20 most intense monoisotopic precursors with intensity > 100 cps, from a candidate mass range of 800-2250 *m/z*, and with charge states 2-6 were selected for MS2 fragmentation with one of four fragmentation regimes: CID, EAD and two EAD methods with supplemental CID fragmentation termed default and low energy EAciD. For all fragmentation methods parameters wer: Zeno trapping, on; Zeno threshold, 100,000; TOF mass range, 50-4500 *m/z*; Dynamic background Subtraction, on; and former candidate ion exclusion, for 6 s after 2 occurrences. EAD simultaneous trapping mode was used for EAD and EAciD methods, the latter having dynamic collision energy enabled. The default equation for dynamic collision energy was employed for CID and default EAciD methods, and a modified equation with half the gradient was used for low energy EAciD. Tuneable parameters for EAD-based methods were as follows: electron beam current, 5000 nA; electron kinetic energym 8 eV; radio frequency, 200 Da; reaction time, 20 ms; and accumulation timem 50 ms. For CID fragmentation, an accumulation time of 50 ms was also used. Injections were made in triplicate, with randomised injection order within replicates for each fragmentation method.

### Glycopeptide Identification

Glycopeptides were identified using Byonic (Protein Metrics, v4.3.4) searching against the human database of high confidence proteins (20,350 proteins, downloaded from UniProt 25/11/2021) appended with porcine trypsin and 199 bovine serum proteins [37], and a glycan database provided by Byonic containing 182 human *N*-glycans and 6 common *O*-glycans (set as “rare 1” modifications). Cleavage was set as fully specific, C-terminal to arginine and lysine, and permitting two missed cleavage events. Propionamide at cysteine was set as a fixed modification, and variable modifications were oxidation at methionine and deamidation of asparagine (set as “common 1” modifications). A total of two common modifications and 1 rare modification were permitted per glycopeptide. A precursor mass tolerance of 20 ppm, and a fragment mass tolerance of 0.1 Da were used. Fragmentation mode was set to HCD for CID, EThcD for EAciD and ECD for EAD methods respectively.

### Statistical Analysis and Data Visualisation

Principal component analysis and heatmaps of glycopeptide peptide spectral matches (GPSMs) and their assigned Byonic scores were generated with ClustVis [38]. Non-linear regression analysis was performed to model the score distributions of GPSMs. Two-tailed unpaired Student’s *t-*tests were performed for comparing the means of two groups and Fisher exact tests were performed to test the significance of association of categorical variables. Two-way ANOVA and Dunnett’s multiple comparison test were performed when comparing quantitative variable changes based on two independent variables. Statistical testing was performed in GraphPad Prism (v 9.4.0).

## Results

### Tuneable EAD Parameters

The major feature distinguishing EAD from other ExD methods is the ability to fine-tune the underlying instrument parameters to suit specific experimental needs. We surveyed several tuneable EAD parameters to optimise conditions for glycopeptide identification. To obtain a sample with a complex mixture of mammalian *O*- and *N*-glycopeptides, we enriched glycopeptides from a tryptic digest of proteins secreted from human A549 cells. We repeatedly measured this sample with LC-MS/MS with EAD fragmentation and assessed the effect on glycopeptide identification by Byonic of individually changing each tuneable EAD parameter: radio frequency (RF), electron beam current, electron kinetic energy (KE), and reaction time.

The ZenoTOF 7600 instrument applies an RF voltage to trap ions in the EAD cell to increase the efficiency of EAD fragmentation. A higher RF can be used to trap product ions across a wider mass range [39]. We tested RF from 100-300 Da and found that using a lower mass cut-off (LMCO) value of 200 Da was optimal for glycopeptide identification and spectral quality (Figure **1A**). Increasing the electron KE increases the thermal energy input to the precursor ions, improving the dissociation of molecules when electron capture alone is insufficient. However, increasing the electron KE too far reduces the cross-sectional area for electron capture, and results in the electrons more frequently inducing vibrational dissociation and reducing c and z-type ion signals [28]. We tested KE from 0-12 eV, and found that higher values 8 or 12 keV performed better than lower values (Figure **1A**). The electron beam current parameter controls the flow of the electrons used to induce fragmentation in EAD. Precursor fragmentation is proportional to the electron beam current, but secondary electron capture events that neutralise fragment ions and render them undetectable by the mass analyser are more frequent with higher electron beam current [40]. We tested electron beam currents from 2500-7500 nA and found 5000 nA was optimal for glycopeptide identification (Figure **1A**). Increasing reaction time also improves the fragmentation efficiency in EAD, although acquisition times must be compatible with LC peak widths to achieve comprehensive analyte coverage. We tested reaction times from 10-45 ms, and found that the longer 20 or 45 ms reaction times were preferrable (Figure **1A**). We subsequently refined the EAD parameters for KE and reaction time, in combination. We once again analysed our complex glycopeptide sample and tested four EAD methods varying combinations of KE (8 and 12 eV) and reaction times (20 and 45 ms). We observed a greater number of unique glycopeptide identifications in the methods with the shorter reaction time of 20 ms (Figure **1B**). With a 20 ms reaction time, we found that adjusting KE did not significantly change the overall number of glycopeptide identifications. However, glycopeptide identifications with a KE of 8 eV tended to have higher scoring GPSMs than the 12 eV method (Figure **1C**). Therefore, we selected an optimized EAD fragmentation method with a KE of 8 eV, reaction time of 20 ms, RF of 200 Da, and electron beam current of 5000 nA for maximum identification of glycopeptides with high quality MS/MS spectra.

**Figure 1.**
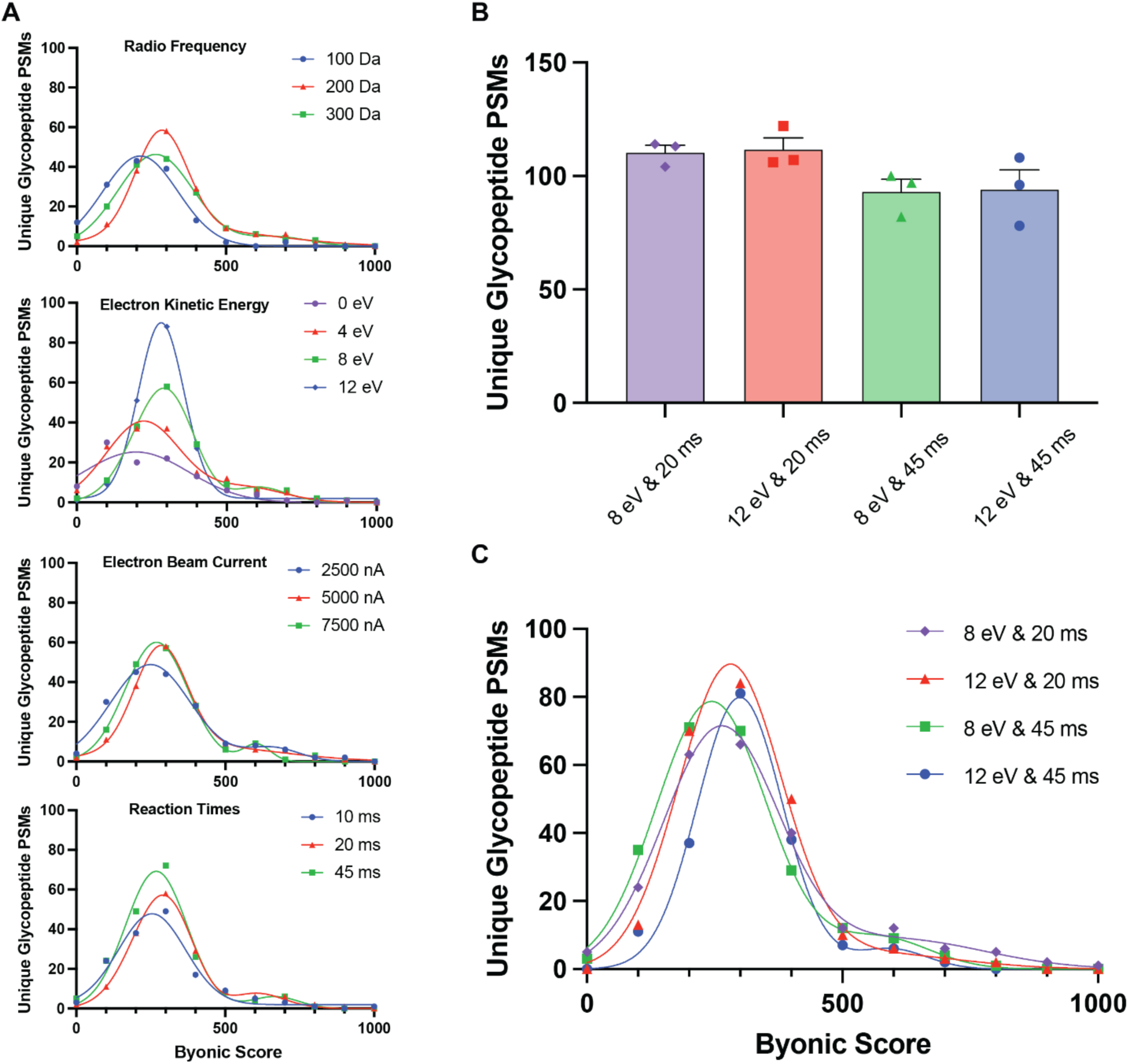
Optimisation of EAD tuneable parameters for glycopeptide characterisation. **(A)** Distributions of Byonic scores of unique glycopeptide PSMs from LC-MS/MS analysis of a complex glycoproteome of enriched glycopeptides from proteins secreted by human A549 cells, varying radio frequency (RF), electron kinetic energy (KE), electron beam current, and reaction time, in the absence of any other changes. **(B)** Mean number of unique glycopeptide PSMs ± SEM (n=3) with the joint modifications of KE and reaction times. **(C)** Distributions of Byonic scores of unique glycopeptide PSMs with the modification of both KE and reaction times. Distributions show PSM counts in bin widths of 50. Non-linear curves were fit to the data using a sum of two gaussian curves model (Prism v9.4.0).

### Comparison of CID and EAD Fragmentation

Having determined optimal EAD parameters for glycopeptide analysis, we next performed a comparative analysis of four fragmentation schemes: standard CID (beam-type), EAD, and two mixed fragmentation methods combining EAD with supplemental CID fragmentation termed hereafter as EAciD. The EAciD methods featured two different levels of supplemental CID, both supplied based on a rolling collision energy (CE) equation determined by precursor ion *m/z* and charge state. The first EAciD method (default EAciD) used a standard rolling CE equation. The second method (low energy EAciD) used an adjusted rolling CE equation with a 50% reduced gradient. We tested these fragmentation schemes on the complex enriched glycopeptide sample originating from proteins secreted from human A549 cells. Across all methods we identified 1,132 human protein groups and 834 unique glycopeptides, including identifications of both *O*-glycopeptides and each of the major glycan classes of *N*-glycopeptides (oligomannose, hybrid and complex). We performed principal component analysis (PCA) of the highest score attributed to each unique GPSMs by Byonic for each method (Figure **2A**). We observed tight clustering between the replicates of each method, and next between EAD and low energy EAciD methods, consistent with the methods providing robust yet distinct glycopeptide identification performance (Figure **2A**).

**Figure 2.**
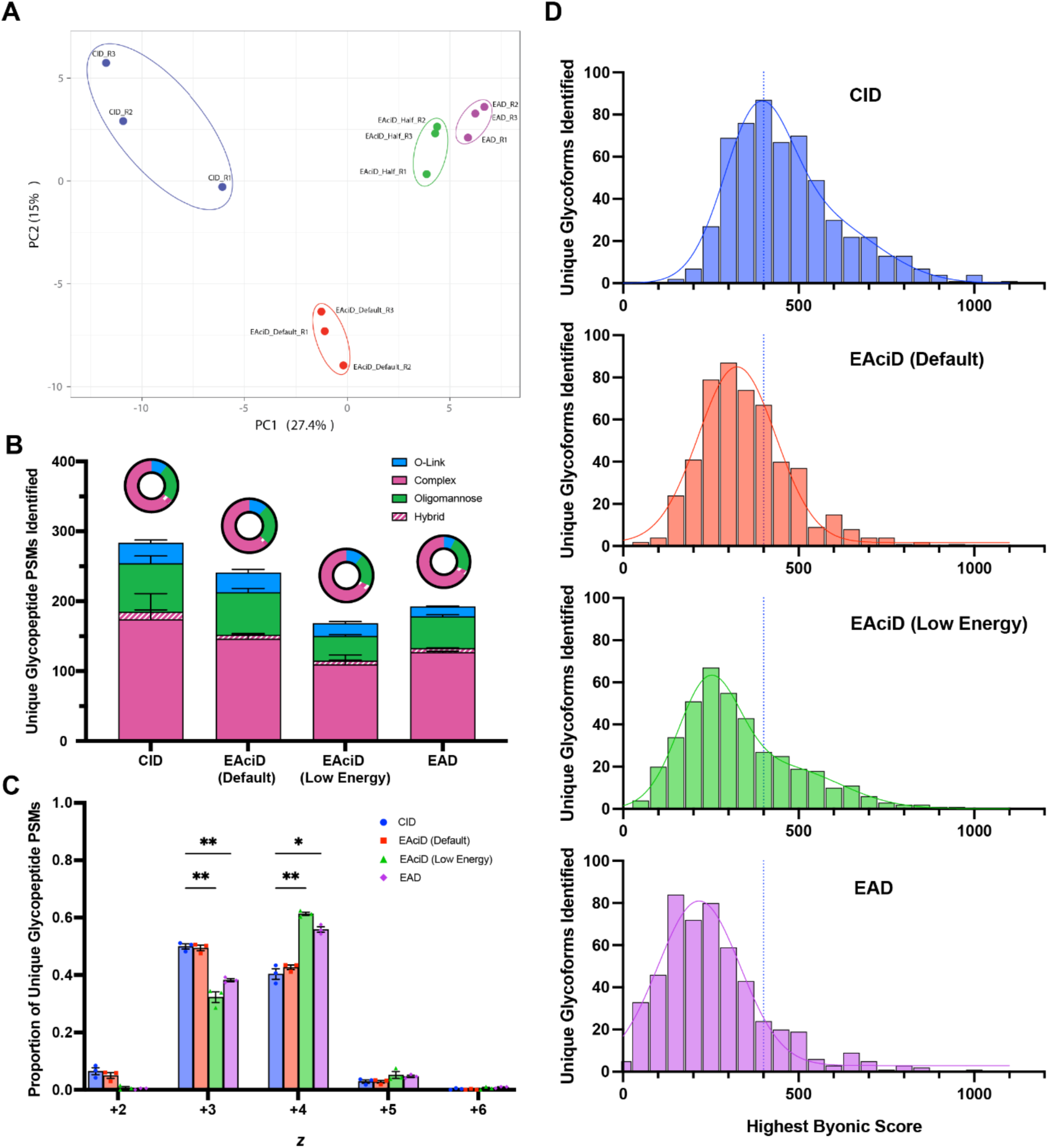
Variance between the performances of CID, EAD and hybrid EAciD fragmentation methods. LC-MS/MS analysis of an enriched glycopeptide sample originating from secreted proteins from human A549 cells (n=3) under four fragmentation schemes: CID (blue), default EAciD (red), low energy EAciD (green) and EAD (purple). **(A)** Principal component analysis (PCA) of uniquely identified glycopeptide PSMs and their highest associated scores (Byonic™, PMI). PCA was generated using the web-based tool ClustVis. **(B)** Number of unique glycopeptide PSMs (score > 200) stratified into four categories (*O*-GalNAc, complex *N*-glycans, oligomannose *N*-glycans and hybrid *N*-glycans). Relative proportions of each category are graphically represented within the donut chart above each bar. **(C)** Proportions of unique glycopeptide PSMs (score > 200) separated by their associated charge state (*z*) from +2-6. A two-way ANOVA analysis and Dunnett’s multiple comparison test was performed to test significance of each method compared with CID for each charge state (Prism v9.4.0). Significant differences were observed between CID and EAciD (Low Energy) methods at +3 (p = 0.0082) and +4 (0.0081) charge states, and between CID and EAD also at +3 (p = 0.0022) and +4 (p = 0.0103). **(D)** Distribution of Byonic scores of unique glycopeptide PSMs. Values show PSM counts in bin widths of 50. Non-linear curves were fit to the data using a sum of two gaussian curves model (Prism v9.4.0). Dotted blue line corresponds with the peak of this curve under CID fragmentation.

To assess the performance of each fragmentation method, we considered a range of quantitative and qualitative metrics. We first considered whether the different fragmentation methods biased the resulting characterisation of the glycoproteome. For instance, previous work has reported reduced efficiency of ExD methods for glycopeptides with large, negatively charged glycans or precursors with lower charge states [27, 41]. We first asked if the various CID/EAD fragmentation methods identified different proportions of *O*-glycopeptides or the three major classes of *N*-glycopeptides (complex, oligomannose, and hybrid). We found that the relative proportions of each glycopeptide class were comparable with each method (Figure **2B**), and hence concluded that the EAD-based fragmentation methods did not bias identification of specific classes of glycopeptide compared to CID. We next tested if the performance of the fragmentation methods was impacted by precursor charge state. This analysis showed that EAD and low energy EAciD methods tended to identify glycopeptides with higher charge states, compared to CID and higher energy EAciD (Figure **2C**). Although a bias for identification of precursors with higher charge states likely contributed to the lower number of identifications by EAD and low energy EAciD, this did not otherwise bias the coverage of the glycoproteome by these methods.

Optimising methods to obtain the most glycopeptide identifications is a key goal in LC-MS/MS glycoproteomics. However, it is also critical to consider overall glycopeptide spectral quality, as higher quality spectra increase the confidence of glycan compositional characterisation and site localisation. We therefore next considered the quantitative and qualitative performances of EAD and EAciD methods compared to standard CID methods by considering the distribution of glycopeptide identifications and their associated spectral scores (Figure **2D**). By this metric, CID outperformed all other methods by yielding the most unique glycopeptide PSMs, that were also on average associated with higher quality spectra. Our optimised EAD method identified comparatively fewer glycopeptides. However, supplementing EAD with CID fragmentation improved the overall quality of glycopeptide spectra, as determined by a shift in the distributions towards higher scores, and in the case of the default EAciD method, also increased the number of glycopeptide identifications (Figure **2D**).

EAD-type fragmentation can be particularly powerful for determining the precise sites of labile modifications in a peptide. The “delta mod” score provided by Byonic is a metric of the confidence of the localisation of peptide modifications, with scores > 10.0 representing a high likelihood that modifications are correctly localised. We tested the performance of the CID, EAD, and EAciD methods as judged by delta mod scores for *O*-glycopeptides, and found that both EAD and low energy EAciD outperformed CID, with a significantly higher proportion of PSMs passing this confidence threshold (Figure **3**).

**Figure 3.**
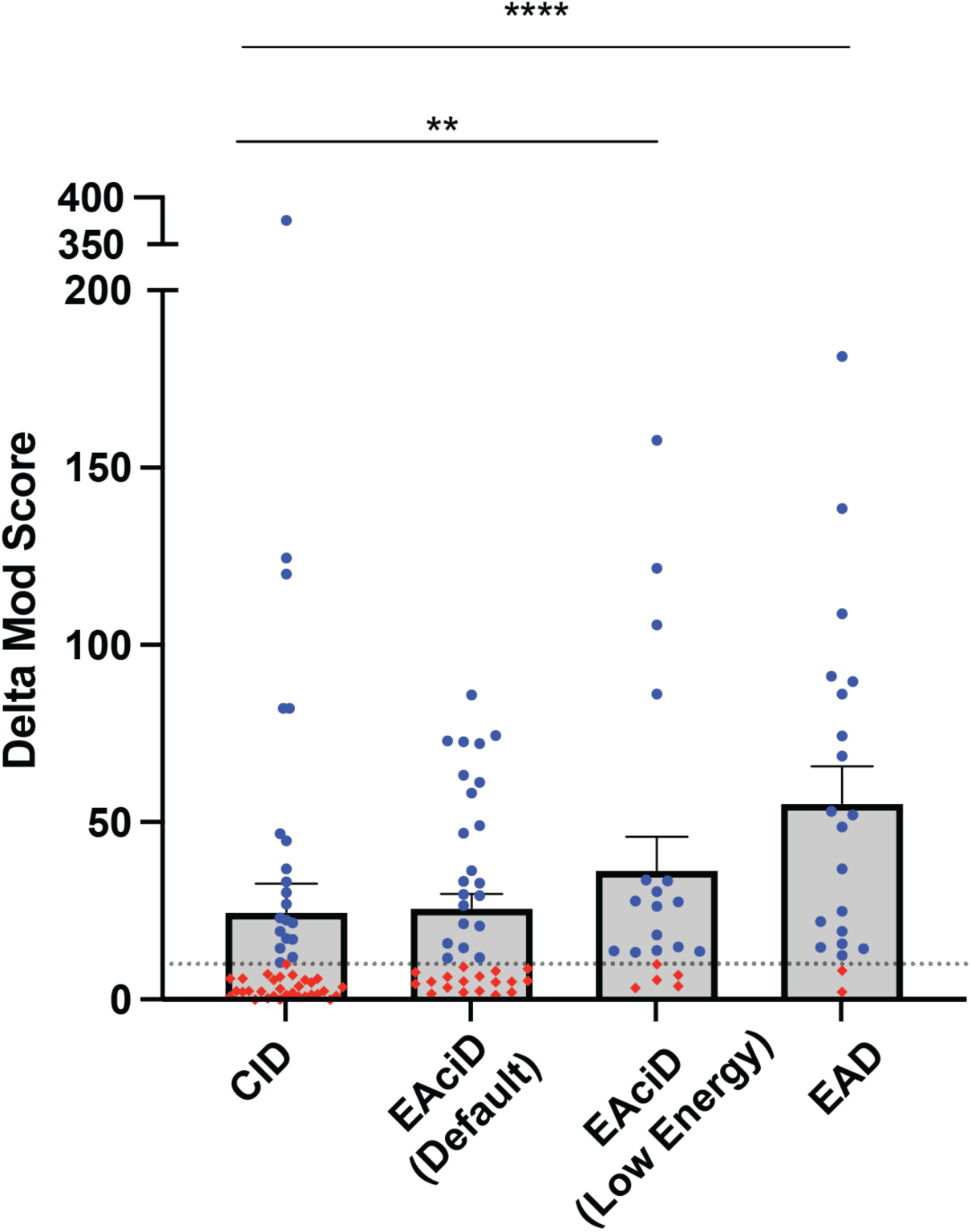
Comparison of CID, EAD, and hybrid EAciD fragmentation methods using a localisation metric. Mean Byonic delta mod score (± SEM) of unique *O*-GalNAc glycopeptide PSMs (score > 200) from LC-MS/MS analysis of an enriched glycopeptide sample originating from secreted proteins from human A549 cells using four fragmentation methods (CID, high energy EAciD, low energy EAciD and EAD). Delta mod scores > 10.0 (in blue) indicate confidence that all peptide modifications are correctly positioned. A Fisher Exact test is performed between CID and EAD-based fragmentation methods based on the number of PSMs that pass and fail the confidence threshold (Prism v9.4.0). A significant increase in the number of PSMs passing the confidence threshold was observed with low energy EAciD (p = 0.0086) and EAD (p < 0.0001).

Different sets of glycopeptides were identified with significantly higher confidence by the CID or EAD-based methods. Of the glycopeptides identified across all methods, 72 had a significantly different score between CID and at least one of the EAD-based methods (Figure **4A**). The majority of these GPSMs (84.7%) had a higher score with CID, consistent with the overall better performance of this fragmentation method. However, some glycopeptides had a higher score with at least one of the EAD-based methods than with CID. We therefore examined these glycopeptides to identify any features of these glycopeptides that correlated with their improved score with EAD. The glycopeptides that were better identified with an EAD method were significantly enriched in complex *N*-glycans (p≤0.01) (Figure **4B**), had significantly higher glycan masses (p≤0.01) (Figure **4C**), and had higher charge states (p≤0.01) (Figure **4D**). In contrast, peptide centric features such as peptide length (Figure **4E**), isoelectric point (Figure **4F**), and peptide mass (Figure **4G**) were not significantly different.

**Figure 4.**
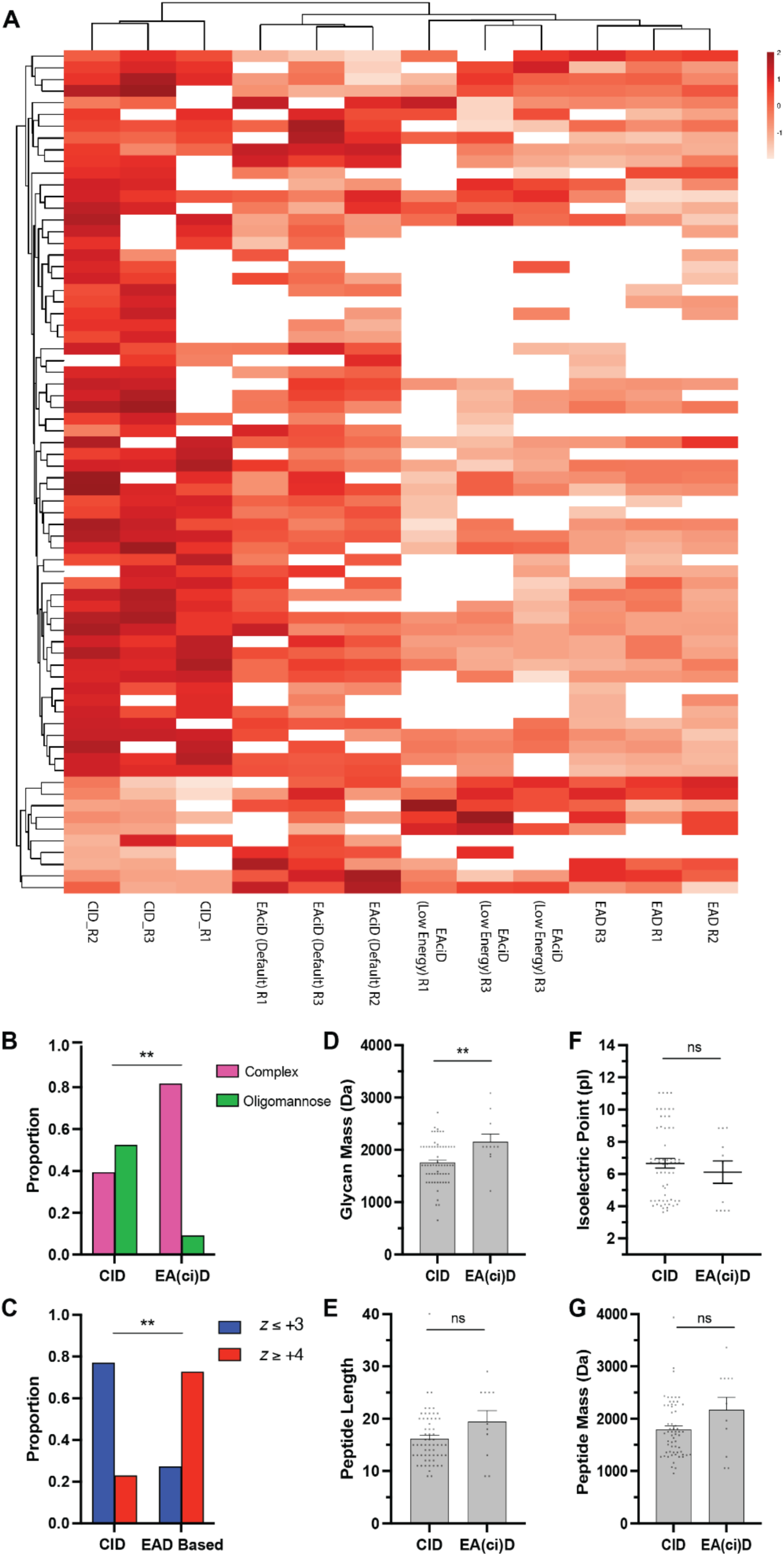
Characterisation of differentially scored glycopeptide PSMs under CID, EAD, and hybrid EAciD fragmentation methods. **(A)** Clustered heatmap of Byonic scores of glycopeptide PSMs (score > 200) from a complex enriched glycopeptide sample from human A549 secreted proteins, analysed by LC-MS/MS with four methods of fragmentation (CID, default EAciD, low energy EAciD and EAD). Scores are represented within the heatmap as their Z-scores, or number of standard deviations from the mean of the entire group. Heatmap clustering was performed using a Euclidean model in ClustVis. (**B-G)** depict information on glycopeptide PSMs that are scored higher in either CID or at least one of the EAD or EAciD methods **(B)** Relative proportion of complex and oligomannose glycans. A Fisher Exact test is performed for statistical significance (p = 0.0069). **(C)** Relative proportion of glycopeptide PSMs with charge states ≤ +3 and ≥ +4. A Fisher Exact test is performed for statistical significance (p = 0.0023). **(D)** Average glycan mass (Da) of glycopeptide PSMs. A two-tailed t-test is performed for statistical significance (p = 0.0055). **(E)** Average peptide length of glycopeptide PSMs. A two-tailed t-test is performed for statistical significance (p = 0.0690). **(F)** The average isoelectric point (pI) of the underlying peptide from each glycopeptide PSM. A two-tailed t-test is performed for statistical significance ( p = 0.471). **(G)** Average peptide mass (Da) of the underlying peptide from each glycopeptide PSM. A two-tailed t-test is performed for statistical significance (p = 0.0549). Statistical tests performed in Prism v9.4.0.

Although spectral scores provide a good indication of overall quality, direct comparisons between the glycopeptide spectra produced by different fragmentation methods can provide additional evidence of the performance of each that may be overlooked by simple metrics. We assessed the performance of each fragmentation method by examining glycopeptide PSMs for different types of glycosylation identified across each dissociation method. Spectra for an Ephrin-A1 (P20827) glycopeptide modified with a HexNAc(2)Hex(8) oligomannose glycan varied by method (Figure **5A-D**). Precursor consumption in EAD and low energy EAciD was low compared with default EAciD and standard CID (Figure **5A-D**). However, the presence of c6 and z6 fragment ions in all three EAD-based methods allowed unambiguous localisation of the *N*-glycan (Figure **5B-D**). Oxonium ions and B- and Y-ions in CID, default EAciD and, to a lesser extent, low energy EAciD allowed characterisation of the glycan (Figure **5A-C**). Spectra for a metalloproteinase inhibitor-1 (P01033) glycopeptide modified with a HexNAc(4)Hex(5)Fuc(1)NeuAc(2) complex glycan revealed both similarities and differences to the oligomannose glycopeptide. CID and EAciD spectra were dominated by oxonium ion and B- and Y-fragment ions useful for glycan characterisation (Figure **6A-C**). The CID spectra featured b- and y-ions that could characterise the peptide backbone, albeit at relatively low intensity compared to oxonium ions (Figure **6A**). Default EAciD spectra for this glycopeptide did not generate intact glycopeptide fragment ions that could be used to localise the glycan (Figure **6B**), but this was possible with low energy EAciD and EAD, which produced informative doubly charged c8++ and z8++ ions (Figure **6C** **& D**). Analysis of an *O-*glycopeptide from cathepsin D (P07339) showed informative oxonium and B- and Y-fragments with CID (Figure **7A-C**), low precursor consumption in EAD and low energy EAciD (Figure **7C** **& D**), and the presence of the c9 fragment ion in all EAD-based methods that enabled site-localisation of the *O*-glycan (Figure **7B-D**).

**Figure 5:**
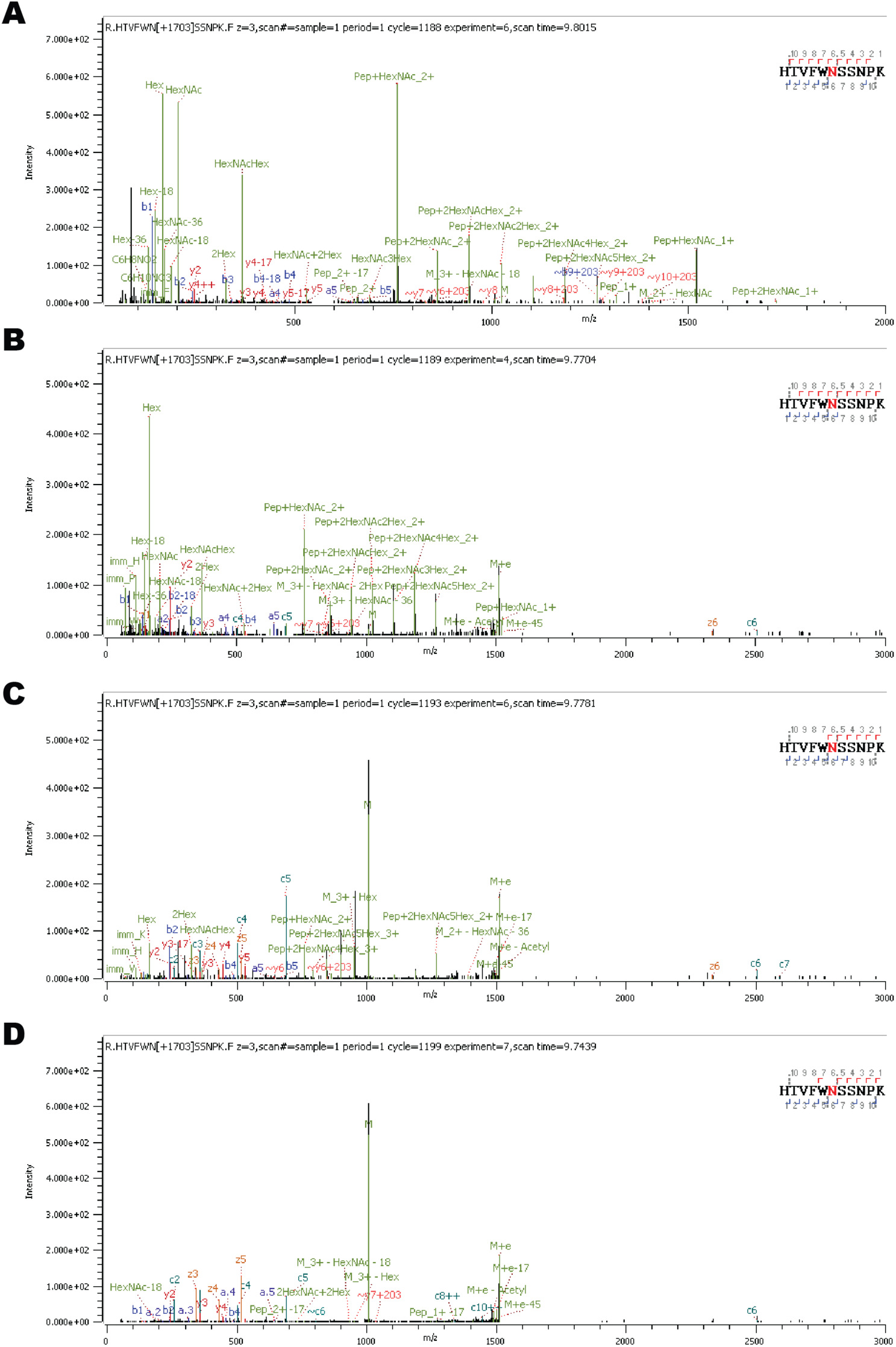
Fragmentation patterns of an oligomannose glycopeptide under various fragmentation methods. MS/MS analysis of a glycopeptide of a human Ephrin-A1 glycoprotein (P20827) with an oligomannose type *N*-glycan HexNAc(2)Hex(8) attached under four fragmentation methods **(A)** CID fragmented (observed 1007.078 *m/z*, 3+ and scoring 554.04) **(B)** default EAciD fragmented (observed 1007.078 *m/z*, 3+ and scoring 398.60) **(C)** low energy EAciD fragmented (observed 1007.083 *m/z*, 3+ and scoring 371.64) and **(D)** EAD fragmented (observed 1007.078 *m/z*, 3+ and scoring 346.22)

**Figure 6:**
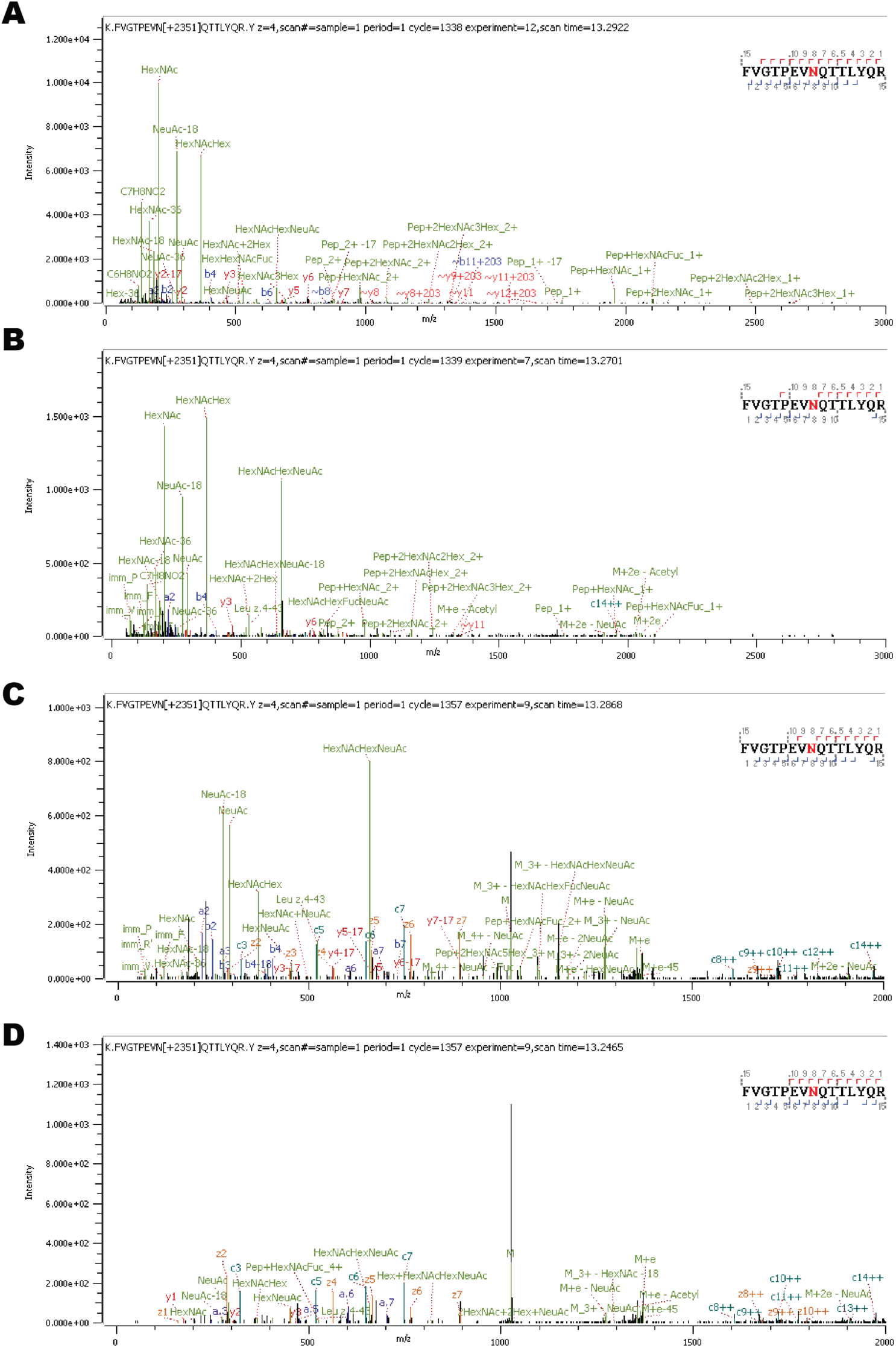
Fragmentation patterns of a complex glycopeptide under various fragmentation methods. MS/MS analysis of a glycopeptide of a human metalloproteinase inhibitor-1 glycoprotein (P01033) with a complex type *N*-glycan HexNAc(4)Hex(5)Fuc(1)NeuAc(2) attached under four fragmentation methods **(A)** CID fragmented (observed 1026.691 *m/z*, 4+ and scoring 677.18) **(B)** default EAciD fragmented (observed 1026.689 *m/z*, 4+ and scoring 494.57) **(C)** low energy EAciD fragmented (observed 1026.681 *m/z*, 4+ and scoring 590.87) and **(D)** EAD fragmented (observed 1026.693 *m/z*, 4+ and scoring 692.49).

**Figure 7:**
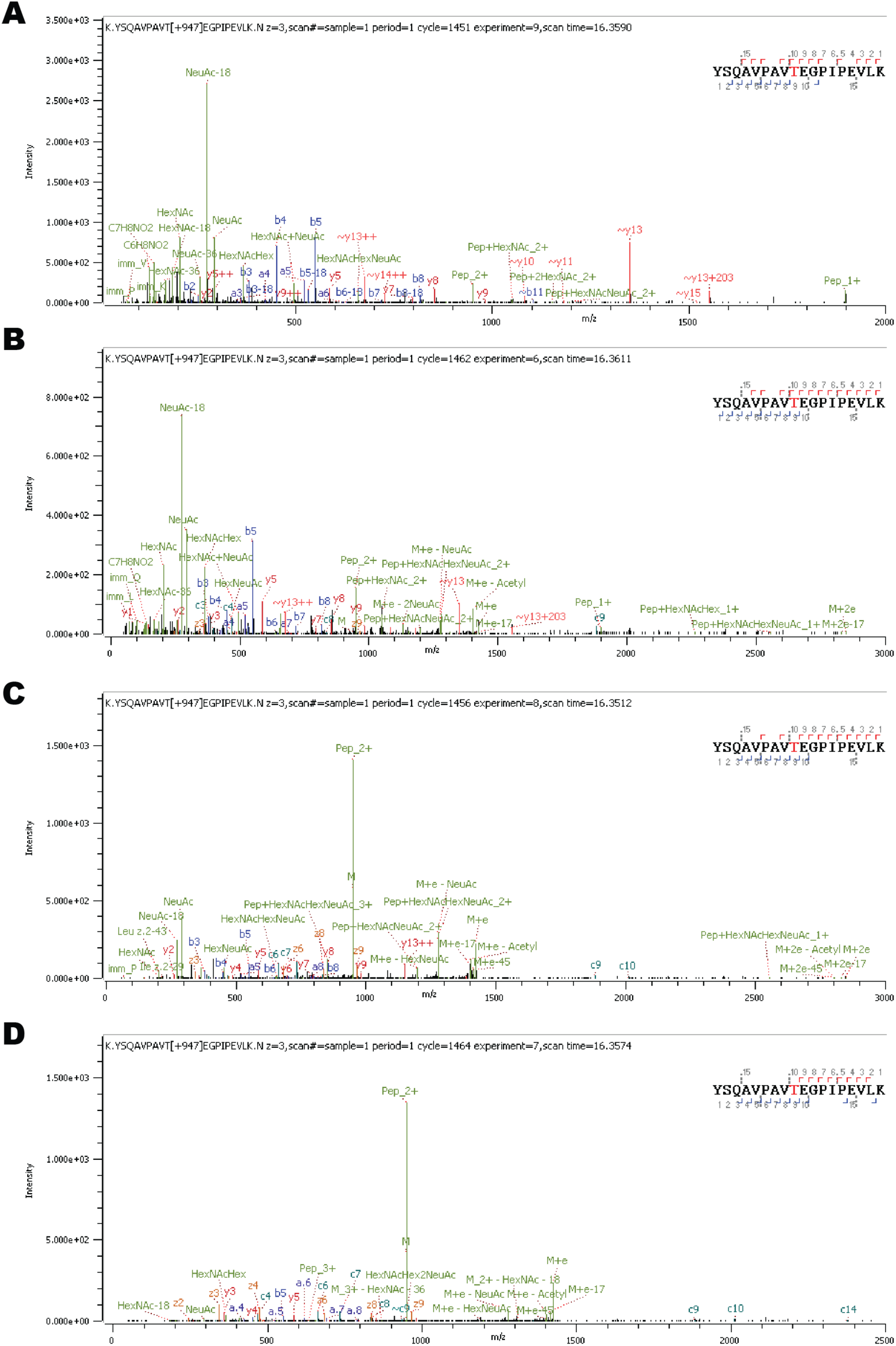
Fragmentation patterns of an *O-*glycopeptide under various fragmentation methods. MS/MS analysis of a glycopeptide of a human cathepsin D glycoprotein (P07339) with an *O-*glycan HexNAc(1)Hex(1)NeuAc(2) attached under four fragmentation methods **(A)** CID fragmented (observed 949.120 *m/z*, 3+ and scoring 889.29) **(B)** default EAciD fragmented (observed 949.122 *m/z*, 3+ and scoring 615.58) **(C)** low energy EAciD fragmented (observed 949.125 *m/z*, 3+ and scoring 482.75) and **(D)** EAD fragmented (observed 949.120 *m/z*, 3+ and scoring 382.25).

## Discussion

Making use of the possibility of tuning EAD parameters on the ZenoTOF 7600, we optimized RF, electron KE current, beam current, and reaction times for analysis of a complex sample of mammalian *O*- and *N*-glycopeptides. We found that the highest number of glycopeptide identifications were observed using EAD with electron KE values of 8 and 12 eV. This is consistent with previous studies that have reported advantages in glycopeptide analysis in the hot ECD range [42], particularly for sialylated glycopeptides [27]. Optimal KE values for EAD fragmentation in glycoproteomic analyses will likely vary depending on the sample glycan composition, although a hot ECD value of ∼8 eV is reasonable for glycopeptide analysis. We found that a moderately increased RF amplitude improved glycopeptide identification, however further increases in RF amplitude began to instead have a detrimental effect. Glycopeptide fragmentation produces ions across a wide mass range, from oxonium ions to large Y-ions, consistent with better performance at moderate to high RF values. Reduced performance at high RF amplitude has been attributed to increased electron energy by RF heating, and loss of ion trapping consistent with our observations of a moderate optimal RF value [35, 43]. We found an electron beam current of 5000 nA was optimal for glycopeptide analysis, consistent with previous reports [40]. Finally, a reaction time of 20 ms was optimal for glycopeptide analysis, consistent with standard methods for peptide analysis. While we identified the optimum parameters for this particular sample, it is possible that other samples, in particular those with different types of glycosylation, may benefit from different EAD parameters.

Directly comparing CID, EAD, and EAciD methods, we found that CID performed best in terms of the total number of glycopeptides identified, while EAD and EAciD were superior in providing localisation information in the form of c and z product ions with an intact glycan. EAciD methods feature peptide backbone fragmentation, c and z fragment ions that can localise the site of glycosylation, and oxonium ion profiles that can support glycan composition assignments, enabling more complete glycopeptide characterisation. However, we found that EAciD methods did come with a cost to the overall number of glycopeptide identifications. We applied two levels of supplemental CID, both based on rolling collision energy. Although both EAciD methods had some beneficial features of CID and EAD, the performance of the two methods was quite different. The default EAciD method performed more like CID, with poorer localisation metrics but increased identifications. In contrast, reducing the supplemental collision energy in the low energy EAciD method improved localisation and generated information rich spectra, but had relatively few glycopeptide identifications. The choice to use CID, EAD, or EAciD for glycoproteomics is dependent on whether the desired experimental objectives favour greater coverage of the glycoproteome or in-depth characterisation of the measured glycopeptides.

## Conclusion

We have developed and demonstrated an optimised workflow for mammalian glycoproteomics using EAD fragmentation on the ZenoTOF 7600. We found that standard beam-type CID could identify more glycopeptides than EAD, but that EAD could provide more information on the precise sites of glycosylation. EAD with supplemental collision energy (EAciD) further improved glycoproteomic analyses compared to standard EAD. This EAciD combination of EAD and CID fragmentation provides highly informative glycopeptide MS/MS spectra suitable for characterising the peptide, glycan, and the site of modification for glycopeptides with diverse *O*- or *N*-glycans.

## Acknowledgements

We thank The University of Queensland, School of Chemistry and Molecular Biosciences Mass Spectrometry Facility for assistance and expertise. This work was supported by a National Health and Medical Research Council (NHMRC) Ideas grant APP1186699 to B.L.S. and C.L.P, and Australian Research Council (ARC) Linkage Infrastructure, Equipment and Facilities grant LE220100068.

